# Who Bears the Burden of Long-Lived Molecular Biology Databases?

**DOI:** 10.1101/552067

**Authors:** Heidi J. Imker

## Abstract

In the early 1990s the life sciences quickly adopted online databases to facilitate wide-spread dissemination and use of scientific data. From 1991, the journal *Nucleic Acids Research* has published an annual Database Issue dedicated to articles describing molecular biology databases. Analysis of these articles reveals a set of long-lived databases which have now remained available for more than 15 years. Given the pervasive challenge of sustaining community resources, these databases provide an opportunity to examine what factors contribute to persistence by addressing two questions 1) which organizations fund these long-lived databases? and 2) which organizations maintain these long-lived databases? Funding and operating organizations for 67 databases were determined through review of Database Issue articles. The results reveal a diverse set of contributing organizations with financial and operational support spread across six categories: academic, consortium/collective, government, industry, philanthropic, and society/association. The majority of databases reported support from more than one funding organization, of which government organizations were most common source of funds. Operational responsibilities were more distributed, with academic organizations serving as the most common hosts. Although overall there is evidence of diversification, the most acknowledged funding and operating organizations contribute to disproportionately large percentages of the long-lived databases investigated here.

## INTRODUCTION

Online databases are critical to research in the life sciences (1). These resources are created to meet a variety of needs, often with the intent to be useful across many areas of biology. For example, they vary in scope and content to address a range of research themes, such that some are dedicated to model organisms (e.g., FlyBase) and others are dedicated to covering a specific biophysical phenomenon (e.g., Signal Recognition Particle Database). As life scientists began to heavily utilize these resources, their impact has become increasingly apparent. One study found that articles describing bioinformatic resources, which includes databases, constitute a third of the most highly cited articles in the scientific literature (2).

As the utility of online databases become clear, new databases began to proliferate. In 1991 the journal *Nucleic Acids Research* (*NAR*) started aggregating articles describing molecular biology databases into an annual issue (3). This issue became known as the “Database Issue” and grew to serve as an important and prestigious venue to announce either new databases or updates to established databases. A recent census of all databases published within *NAR* Database Issues revealed over 1700 unique databases debuted since 1991, and the census was released as an openly available dataset (4). While rapid proliferation of databases indicates a vibrant community, it also illuminates the challenge of sustaining these resources over time. Others have noted that initial funding to create online databases is easier to obtain than funding to maintain those resources (5). Indeed, even highly successful databases, including several databases included in this study, have habitually struggled to sustain themselves over the long-term, especially when funding from government sources wanes (6). For example, the OMIM: Online Mendelian Inheritance in Man, a 50 year old resource, now actively solicits donations. As the website explains, “At the request of the NIH and to ensure long-term funding for the OMIM project, we must diversify our revenue stream.” In another example, the enormously popular KEGG: Kyoto Encyclopedia of Genes and Genomes, which amassed >15,000 citations by the end of 2016, was forced to transition the resource from a “fully government-funded project to a more community-supported project.” As of 2017, only 10% of operational costs are covered by public funding, yet it helps keep the database going.^1^ The pressure of this issue is so intractable for individual databases, an effort to establish a global coalition to sustain core data resources in the life sciences is underway (7, 8).

As online databases of all ages are continually challenged to develop sustainability models, a closer look at long-lived databases is warranted. Databases that have weathered the trials of erratic funding, staffing turnover, data updates, and technology refreshes provide an opportunity to observe the realities of persistence. Long-lived databases are often associated with large, established government or intergovernmental organizations, e.g., the European Bioinformatics Institute in the UK, the Swiss Bioinformatics Institute, or the National Center for Biotechnology Information in the US. Although such organizations have even been referred to as “database juggernauts,” Galperin et al. (9) pointed out that databases are not exclusive to these organizations. Yet we have little empirical evidence of the distribution of contributions in practice. The *NAR* database census revealed a broad set of molecular biology databases that have remained available for > 15 years. This paper discusses the analysis of those databases and addresses two of fundamental questions 1) which organizations fund these long-lived databases? and 2) which organizations maintain these long-lived databases? Given the difficulty of obtaining on-going support relative to obtaining initial support, looking at the distribution of responsibility well into a database’s lifecycle provides a view not reported elsewhere.

## METHODS

### Data Collection

A set of long-lived databases for study was initially identified during creation of a census of molecular biology databases described in *NAR* Database Issue articles between 1991-2016. In that work, 105 databases were identified as having now remained available for >15 years. Not all databases are published in *NAR* Database Issue articles and those that are may not necessarily appear soon after a database’s initial launch; thus, this set is not inclusive of all molecular biology databases available >15 years. However, *NAR* Database Issue articles provide a measure of standardization in terms of documentation and reporting and also serves as a mechanism to collect a broad set of databases without the bias of selecting persistent databases based on relative fame or user-community size. The set of 105 long-lived databases was extracted from the openly available dataset for the *NAR* census (10), and current availability was reconfirmed in March 2018 by accessing each database through associated URLs. Since the time the census was carried out in early 2017, 5 database URLs were now no longer functional and while URLs were functional for another 6 databases, the websites revealed either broken functionality that precluded access to the data or contained discontinued notices recommending the resource no longer be used. Finally, 1 database was formally discontinued, and although data for this resource are archived on a file severer which remains functional, this database was also excluded. Thus, of the 105 databases initially considered for inclusion in this study, 12 were removed as no longer extant, leaving 93 for additional consideration.

The content and structure of database websites is highly variable, and *NAR* Database Issue articles were found to be more consistent sources of funding information. Databases are initially published in *NAR* Database Issues in a “debut” article, and subsequent “update” articles may be published in future *NAR* Database Issues. In order to use articles to determine funding sources that contributed to database *persistence* rather than initial *creation*, only databases with at least one subsequent “update” article were considered and this article had to be published a minimum of five years after publication of the initial “debut” article. These criteria excluded an additional 26 databases for 67 databases in the final analysis set.

### Capture of Contributing Organizations

For the remaining 67 databases, each database’s most recent *NAR* Database Issue update article published prior to 2017 was accessed on the journal’s website at Oxford University Press. Since a database’s most recent update article may have been published prior to widespread indexing of funder metadata circa 2009, the full text was manually reviewed to capture funding organizations. Following the heuristics of Grassano et al. (11), organizations had to be explicitly named within the article to be included and were not inferred from other context. Organizations were only counted once per database, regardless if mentioned multiple times within statements (e.g., for distinct grants). While most funding organizations could be interpreted as providing direct financial support to the databases in the form of research grants, other indirect financial support was reported and is included here. This includes support provided through various types of fellowships or computational resources supplied through academic research centers. Exclusions included gratitude to specific people for support that can ostensibly be considered “moral.”

Both the text within the *NAR* article and information found on the database websites were used to determine the organization that currently takes primary responsibility for hosting and operating the database. Specifically, author affiliations, URL domain names, website branding, and “about” pages were reviewed to identify operational homes. Both the country of the operating organization (as ISO 3166-1 alpha-3) and the organization name were recorded. In most cases, this could be resolved to a single organization in a single country. Exceptions include two databases run by consortia with an international composition, and here a code of “INT” was used.

Although sub-organizations were often reported for both funding or operation, e.g., the National Library of Medicine for the US National Institutes of Health or the Interdisciplinary Nanoscience Center at Aarhus University, this granularity was inconsistent, a complication noted in previous efforts to evaluate research funding (11, 12). Thus, the highest level of the organization is reported here. Once complied, organization names were checked for standardization and classified. One classification strategy is binary assignment as “public” or “private,” but more variability was observed here and organizations could not be cleanly assigned using this strategy. Conversely, more granular funding model classifications (e.g., 13–15) were largely too detailed to fully apply in practice here. Thus, for the purposes of this study, organizations were classified as “government” if situated within a governing body of a nation or nations (e.g., Spanish Ministry of Economy Industry and Competitiveness), as “academic” if the organization’s main purpose is higher education (e.g., Kyoto University), as “industry” if the main purpose is to function as a commercial business (e.g., New England Biolabs), as “philanthropic” if the main purpose is to function as a charity or non-profit (e.g., Wellcome Trust), as a “consortium/collective” if comprised of a group of organizations formally working together towards a common scientific purpose (e.g., WormBase Consortium), and as a “society/association” if primarily serving as a professional or learned organization with individual memberships (e.g., Belgian Society of Human Genetics). One challenging classification is the European Molecular Biology Laboratory, which was classified as “government” since it is an intergovernmental organization. In this case, funding is provided by 22 member states and not from the government of a single nation.

### Data Analysis and Availability

With operating and funding organizations identified for each database, the statistical programming language R 3.3.3 to was used to obtain aggregate descriptive statistics. The data were transformed using the RStudio environment (1.0.136) and the package dplyr_0.7.4 from the tidyverse “core,” attached via the tidyverse_1.1.1 “meta” package (16). Analyses were performed using base functions in R and the package moments (17) and visualized using the package ggplot2_2.2.1 (18). The explicit steps of analysis are detailed in the documentation provided with the R scripts and data files in the associated openly available dataset for this work at https://doi.org/10.13012/B2IDB-3993338_V1. Additionally, working copies of data and scripts are provided on GitHub at https://github.com/1heidi/nar_persistent.

## FINDINGS

### Financial Support

Through review of acknowledgments and funding statements within articles, funders were attributed for 63 (94.0%) of the 67 databases. This percentage is on pare with evaluation of life science journals, where results show ~90% of articles report funders (11). A total of 89 unique funding organizations were identified 176 times (supplemental Table 1). The distribution of the number of databases supported by each unique funding organization was non-normal and ranged from 1 database to 29 databases (*M* = 2.0, SD = 3.4 with skewness of 6.1 and kurtosis of 45.6). Correspondingly, distribution of the number of unique funding organizations attributed within each database article was non-normal and ranged from 1 funder to 20 funders (*M* = 2.8, SD = 2.8 with skewness of 3.9 and kurtosis of 22.8).

To examine the diversity of funding sources, the 89 individual funding organizations were mapped to categories. The results revealed acknowledgement of 29 government, 19 industry, 13 academic, 12 philanthropic, 9 society/association, and 7 consortium/collective organizations. The funding codes were aggregated for each database, and since 28 unique combinations resulted (supplemental Table 2), codes were condensed where multiple funders from the same category were reported. For example, the AAIndex: Amino Acid Index Database included funding from the Ministry of Education Culture Sports Science and Technology of Japan (G), Japan Science and Technology Agency (G), Kyoto University (A), and University of Tokyo (A), which was condensed to AG, representing the two different categories of funders reported.

Analysis revealed that the majority of the databases reported a single funding type (n = 38; Figure 1, first panel). Dependence solely on government funding organizations was the most common category (n = 34), making up 54.0% of the 63 databases with funding reported. Only 4 databases (6.3%) did not attribute any funding to any government organizations. The other types of organizations tended to offer support in combination with other sources. The 13 unique academic organizations were attributed in 15 databases (23.8%) yet only a single database relied solely on academic organizations. Similarly, the 12 unique philanthropies were acknowledged for 12 (19.0%) of the databases, only 2 of which reported sole funding by philanthropies. Organizations from industry were the second most numerous category with 19 unique organizations reported, yet 11 of the 19 contributed to a single database, EMBL-EBI’s IMGT/HLA. The remaining 8 industry organizations were mentioned for 7 other databases. Finally, the 7 unique consortium/collective organizations supported 6 (9.5%) databases, and the 9 unique society/association organizations were reported as funding 4 (6.3%) of the databases.

**Figure 1.**
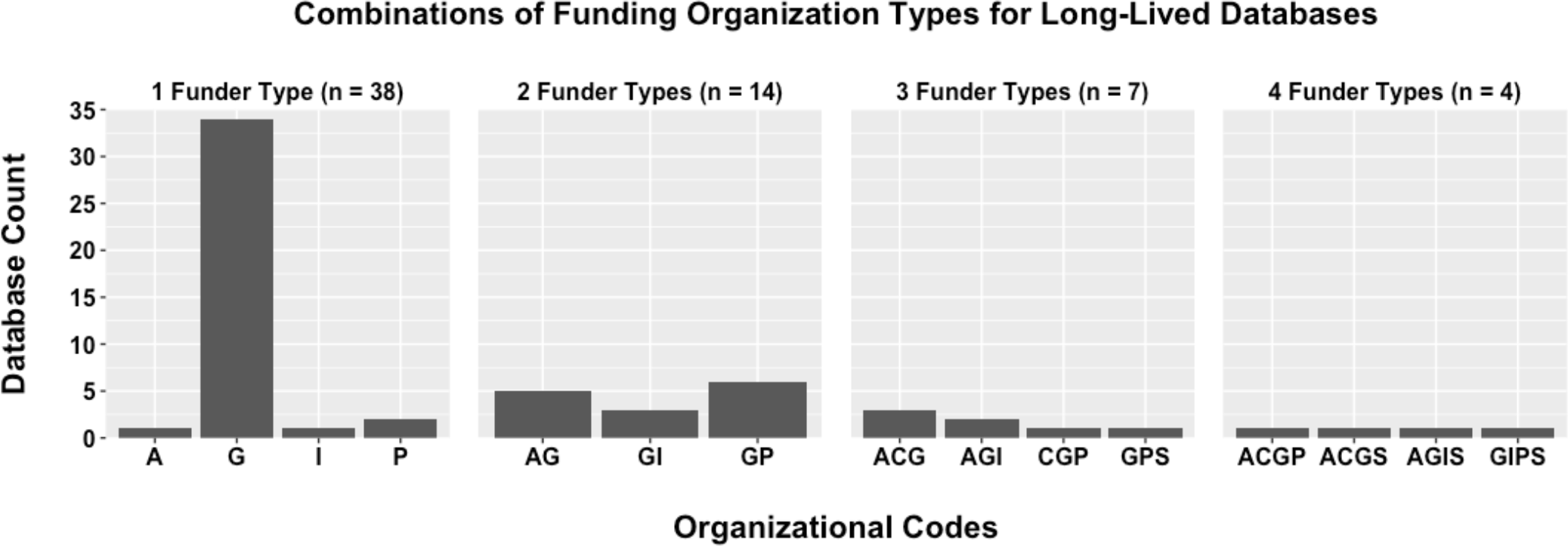
The distribution of funding organizations for the 63 reporting databases shows none attributed more than four distinct types of funders, with the majority of databases attributing support to only government organizations. Organizational codes are A (academic), C (consortium/collective), G (government), I (industry), P (philanthropic), and S (society/association).

The above describes distribution of the types of funding organizations that support these long-lived databases overall, but some databases did not just report one type of funding organization. Instead, they reported only one specific organization altogether. Examination of those that reported a single organization is of interest since it indicates a potentially risky level of dependence on that sole organization. For the 34 databases that reported funding exclusively from government organizations, 20 databases reported a sole government funding source with 14 reporting support from two or more distinct government organizations (supplemental Table 2). Other organizations were less likely to provide sole support with only 1 database reporting a single academic funder, 2 reporting a single philanthropic funder, and 1 reporting a single industry funder. Thus, of the 63 databases with funders reported, 24 (38.1%) appeared to depend on a single funding source only, leaving a majority of 61.9% drawing from multiple funding streams. For comparison, the Organisation for Economic Co-operation and Development (OECD) recently conducted 47 interviews with managers of data repositories which varied in both age and domain. In their results, 24 (52%) reported funding from more than one source (19). Likewise, this finding empirically supports the observation by Gabella et al. that funding streams are often diversified (15).

Government organizations dominate the distribution of funding organizations contributing to the persistence of molecular biology databases and wholly comprise the top 10% of most reported funders as presented in Table 1. While several US organizations are implicated in support of a high number of databases, organizations from the UK, Japan, and Denmark all are represented as individual countries. Two organizations of international status, the European Commission and the European Molecular Biology Laboratory, are also included.

**Table 1.**
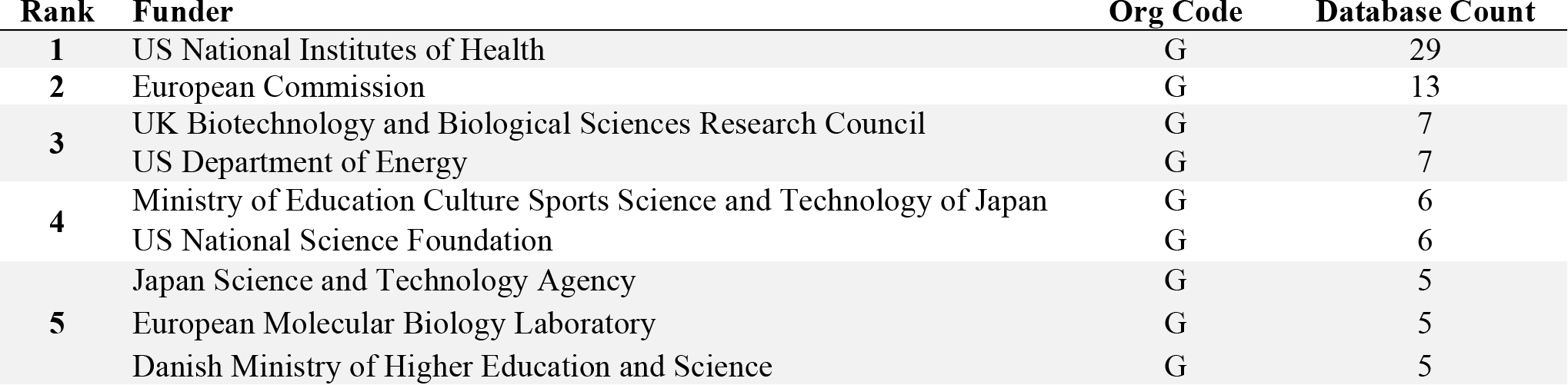
Top 10% of organizations most frequently reported as funding long-lived molecular biology databases.

### Operational Support

A total of 50 unique operating organizations were identified (supplemental Table 3). The distribution of the number of unique databases supported by each operating organization was non-normal and ranged from 1 database to 8 databases (*M* = 1.4, SD = 1.2 with skewness of 4.0 and kurtosis of 20.4). When organizations that operate the 67 databases were assessed, responsibilities were found to be spread across 13 unique countries (Figure 2, Panel A). Two databases were categorized as international (“INT”) based on split of responsibilities born by consortia that work across distinct countries. The distribution of the number of unique databases operated within each country was again non-normal and ranged from 1 database to 26 databases (*M* = 4.7, SD = 6.9 with skewness of 2.3 and kurtosis of 7.7). A total of 46 (68.7%) of the 67 long-lived databases are operated within the US, UK, and Japan.

**Figure 2.**
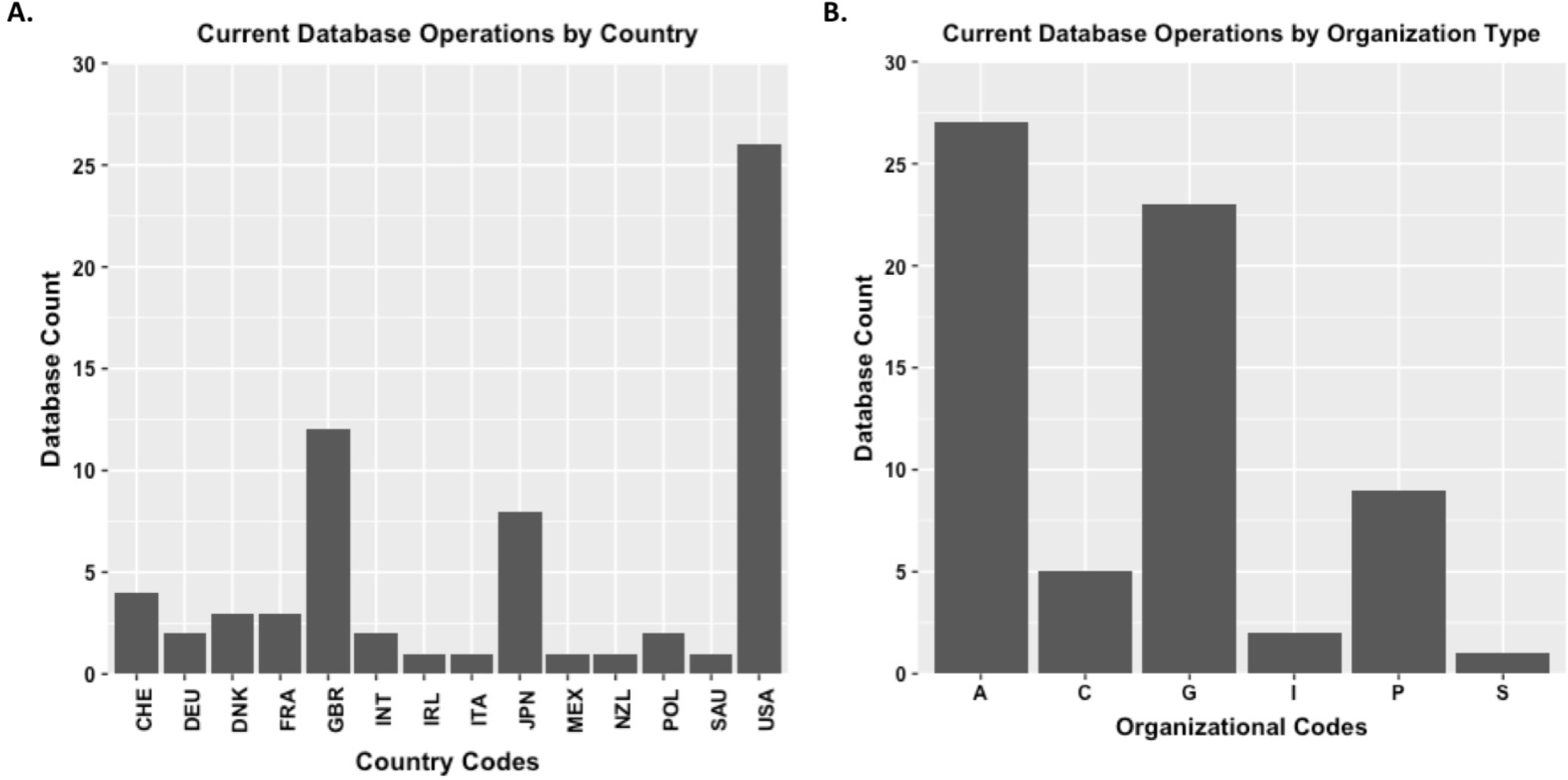
Panel A: Operational responsibilities for 67 long-lived databases are spread across 13 distinct countries. Country codes provided follow the ISO standard, with the exception of “INT” for “international” when operational responsibilities were determined to be split across multiple countries. Panel B: Operational responsibilities for 67 long-lived databases were found to be most likely to fall to academic and government organizations. Organizational codes are A (academic), C (consortium/collective), G (government), I (industry), P (philanthropic), and S (society/association).

As for funding organizations, the operators were analyzed for diversity of organizational type. The 50 individual operating organizations were mapped to the same categories, resulting in identification of 28 academic, 10 government, 5 philanthropic, 4 consortium/collective, 2 industry, and 1 society/association organizations. Operational responsibilities for two databases appeared split across multiple organizations; however, in both cases all organizations were universities. Thus, a clean association of all 67 databases to a single operating organization type was possible. The results are show in Figure 2 (Panel B).

Analysis reveals that the 28 individual academic organizations operated 27 unique databases while the 10 individual government organizations operate 23 unique databases. The 5 philanthropies operate 9 databases, 4 consortium/collective organizations operate 5 databases, 2 industry organizations operate 2 databases and finally the 1 society/association organization operates the remaining database. Interestingly, the distribution of types of organizations that operate databases is more diverse than those that fund databases. Table 2 below shows that 2 organizations categorized as philanthropies, Kazusa DNA Research Institute in Japan and the Jackson Laboratory in the US, operate a total of 6 long-lived databases, or nearly 10% of the entire set analyzed here. The government organizations within the top 10%, namely the European Molecular Biology Laboratory (as an intergovernmental organization), the Swiss Institute of Bioinformatics, and the US National Institutes of Health, operate 16 databases, representing nearly a quarter of long-lived molecular biology databases.

**Table 2.**
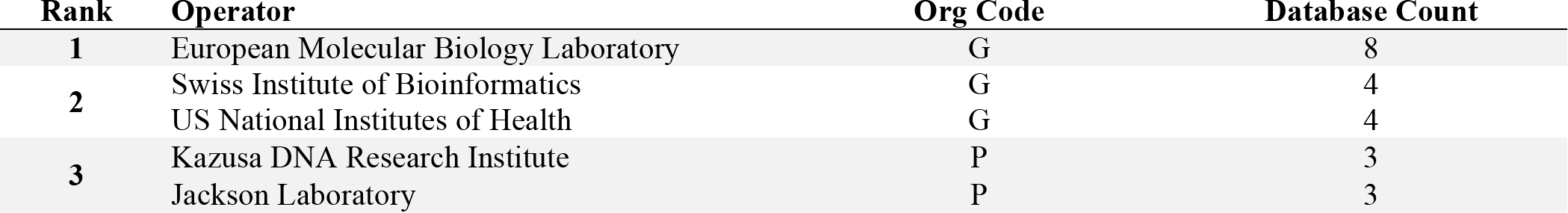
Top 10% of organizations most frequently operating long-lived molecular biology databases.

## DISCUSSION

### Who Bears the Burden?

Perpetual funding for resources that broadly serve scientific communities is a topic of intense interest—and concern (14, 20). Because of research’s contribution to the public good, government organizations often provide research funding in nations across the world. Yet this funding fluctuates. For example, research in Japan and the US, two countries seemingly capable of enduring support in this study, have both weathered challenging budget climates in recent years. Japan’s flat funding and increasing emphasis on connections to industry concerns those conducting research in the basic sciences.^2^ According to the Federation of American Societies for Experimental Biology, the US National Institutes of Health “lost 22% of its capacity to fund research due to budget cuts, sequestration, and inflationary losses” between 2003 to 2015.^3^ Japan and the US are by no means alone in this conundrum; for example, economic circumstances have also negatively impacted Ireland’s research funding (14). Resources that serve research communities, of which databases are a prime example, pose a particularly challenging problem during times of flat or reduced budgets. Databases inherently require on-going support. While some activities may be temporarily postponed without irrevocable damage, such as feature upgrades, failure to attend to technical requirements can quickly render a database entirely non-functional. Notwithstanding their willingness to provide initial funding, government organizations simply do not have large enough budgets to support all databases in perpetuity (21), and many high-profile examples of government organizations withdrawing continued support exist (22, 23). Database providers are urged to “think about new ways to do business” and identify alternative funding sources (6). Without the security of stable government funding, diversified funding streams are desired to avoid disruption of database availability and quality, and as such, diversification is considered a key element in database sustainability (19).

Despite the potential precariousness of government funding, this study shows government organizations remained a source of funding for molecular biology databases long into their lifetimes. Indeed, the vast majority of databases evaluated here continued to depend on some government support. However, a slim majority (54.0%) relied solely on government organizations and an even smaller percentage (31.7%) relied on a single government source. Thus, the data show many of these long-lived databases do, in fact, draw on diverse funding streams. Alternatives to funding from government organizations fell into a number of categories. Academic organizations, namely universities, and private philanthropies both contributed funding to approximately 20% of the databases. Industry organizations were numerous in number; however, the majority were attributed to a funding statement for single database such that this example may be an outlier. All told, although evidence of diversification was observed, of the 176 mentions of funding organizations within database articles, 83 (47.2%) are attributed to the top 9 funders represented in Table 1, meaning 10% of funding organizations identified here make up nearly half of all funding attributions.

The organizations that provide operational support are smaller in number, but more distributed across the organizational types. Here, academic organizations were more likely to bear the load by providing operational support for long-lived databases. Attwood et al. (24) noted a trend of responsibilities falling to institutions, and concluded future support from institutions will be “key” for the sustainability of databases. Within the study here, no one academic institution was found to host more than two databases; i.e. individual universities seem unlikely to assume responsibility for multiple long-lived molecular biology databases. This is unsurprising given the challenge academic organizations face in their attempt to equitably support research infrastructure across all domains, including the Social Sciences and Arts and Humanities, and not just Science, Technology, Engineering, and Mathematics (STEM) fields. On one hand, the large number of academic organizations providing operational support signals further diversification. However, similar to funding, there is also a concentration of operational responsibilities falling to a small number of unique organizations, namely government and philanthropic organizations. The results indicate that the top 10% of operating organizations bear the responsibility for a third of the databases studied here. While concentration into a few organizations carries the benefit of shared infrastructure, a shift in stability or priorities for any of these organizations could likewise jeopardize the operational support currently provided.

### Redistribution of Burden

Alternative funding models are often discussed in the context of database sustainability, and many of the long-lived databases within this study have faced economic challenges. In 2016, the National Institutes of Health’s National Human Genome Research Institute told 9 databases they must identify new funding models by 2020 (6); 6 of those 9 are included in this study. Alternative funding models often include targeting philanthropies, a model already in place for several of the persistent databases analyzed here. Both the Bill and Melinda Gates Foundation and the Institute Pasteur are absent despite being prominent philanthropic supporters of health research; however, the first and second most prominent philanthropies, the Wellcome Trust and the Howard Hughes Medical Institute, respectively, were both identified here as contributors to these long-lived molecular biology databases (12).

The work herein identified streams of support less frequently discussed, namely industry, consortia/collectives, and societies/associations. Organizations that fall into the consortia/collective and societies/associations categories often have limited income sources themselves depending on size and scope. However, their contributions to the longevity of these databases should not be disregarded, and additional avenues here could be examined. Likewise, given the potential capacity, industry strikes one as underutilized. However, Berman and Cerf (25) commented on a notable lack of incentives for industry to participate in sustaining research infrastructure that serves public good.

A more often discussed strategy is adoption of various fee-based models, including deposits fees or access fees. However, both options are controversial. Fee-based strategies rely on users’ willingness and capacity to pay and is likely to disadvantage those without adequate access to funds. Furthermore, as Oliver et al., (26) points out, the source of payment is still likely to originate from grant funding, albeit indirectly and with the added inefficiency of institutional costs included. Additionally, the expectation of free availability of data as part of the open science movement has created its own conundrum as public funding expended to *create* research data gets conflated with public funding (or lack thereof) to *preserve* research data. The OMIM: Online Mendelian Inheritance in Man is attempting to circumvent these issues to establish a user-based funding stream, but through actively soliciting donations from database visitors instead of imposing fees. Notably, OMIM was the only database which explicitly included self-funding in its *NAR* article. Since funding statement requirements have largely been enforced through mandates from funding organizations, self-reference may not be a common practice as of yet. However, disclosure will be necessary in the future as more databases are challenged to pay their own way.

### Limitations

Determination of funding sources is a highly challenging endeavor (12). Funding statements are not always available. Even when they are present, the context in which authors reference funding must be interpreted (11). While it is fortunate that funders could be attributed to a high percentage of databases covered here, the data should not be over-interpreted. In no case was the amount of support reflected in the funding statement, and the distribution will not be equal across all organizations mentioned. Thus, while this study shows which organizations contribute to funding long-lived databases, it does not reveal the relative or absolute extent of those contributions. As Kirby notes, it is not possible to weigh contributions without knowing more information, including the respective monitory allocations from all grants acknowledged (27). Furthermore, all parties that contribute funding are not necessarily attributed within articles. For example, KEGG’s funding statement attributed government sources only, but in actuality funding comes from multiple organizational types (M. Kanehisa, personal communication, June 30, 2018). Grassano et al. also found that authors often do not attribute their employer as providing funding (11). Finally, the results here identified organizations explicitly named within funding statements, but the origin of an organization’s own resources can also be unclear. For example, a university may be drawing funds from a myriad of sources, including government, philanthropic, or industry; yet all that can be determined is that a given university was acknowledged in the article. A more detailed understanding requires careful articulation and accounting of all activities and funding sources that contribute to sustaining a long-lived database, including activities nominally thought of as “in-kind.” An effort to tease apart this accounting by working directly with the database providers may serve as an exciting, albeit highly challenging, opportunity for follow-up. In the meantime, the work herein provides a first look into which organizations are acknowledged as providing any funding contribution for these long-lived databases, regardless of magnitude.

While these databases have been categorized as long-lived based on current availability of the database, it is important to note that each may not be under active development and the reported funding found here may have lapsed already. Indeed, on inspection some databases do appear to be in a static state. This may be considered a limitation of this study; however, even a static database requires on-going attention (and therefore support) to ensure routine care, such as rebooting servers or making sure URLs remain operational. Furthermore, caution is necessary when attempting to assess database status. For example, the REBASE website has a vintage interface but bears ample evidence of attentive upkeep when examined more carefully. Finally, it is not clear if static databases are, in fact, a feature of persistence. One can imagine ways in which the ability to sustain periods of no or low development is beneficial as funding ebbs and flows. Taken to the extreme, purposefully minimizing the need for extensive further development by explicitly building resources that rely on relatively static reference data could be a useful strategy. Investigating these potential strategies is a topic ripe for future investigation as factors that contribute to persistence are investigated further.

## CONCLUSIONS

This study sought to identify which organizations support long-lived molecular biology databases. Through assessment of funding statements for the databases’ *NAR* Database Issue articles, government organizations were found to remain a bedrock of support well into a database’s lifecycle. However, the results also revealed the majority of these long-lived databases do rely on a diversity of funders, and many organizations from other sectors make contributions. In addition to financial support, nongovernment organizations, and especially academic organizations, also play an important part in database longevity by providing operational support. Although signs of diversity are observed, significant portions of these long-lived databases are tied to a relatively small number of organizations for both financial and operational contributions. As data and data-intensive resources, like online databases, continue to proliferate, adequately and equitably supporting them will be a continual strain. While databases as long-standing as those studied here would not be thought of as burdensome in a world of infinite resources, everything is questioned when resources are inescapably limited. The collective value of such community resources seems unequivocal, but it is the current distribution of responsibility—or perhaps more accurately, the lack of feasible mechanisms to redistribute responsibility—that creates strain. As many struggle to reconcile reality with the need for sustainability, this work provide an empirical evaluation of organizations that contribute to persistent databases in practice.

## ACKNOWLEDGMENTS

The author is grateful to Miho Funamori for advice on classification of Japanese organizations, and Ashley Hetrick and Chuck Cook for thoughtful review of the manuscript draft. The author is also grateful to Robert Olendorf and Hoa Luong from the Data Curation Network (http://datacurationnetwork.org) for careful curation of the dataset as well as independent verification that the code executed as expected.

## DATA AVAILABILITY

The data, scripts, and associated documentation for this article are archived and freely available at https://doi.org/10.13012/B2IDB-3993338_V1. The author encourages corrections, reanalysis, reuse, and updating. Active copies of the materials are also accessible on GitHub at (https://github.com/1heidi/nar_persistent) and may be updated since the time of this publication.

## Footnotes

1 https://web.archive.org/web/20170914150208/http://www.kegg.jp/kegg/docs/plea.html

2 https://web.archive.org/web/20171017210413/https://www.nature.com/news/japanese-research-leaders-warn-about-national-science-decline-1.22847

3 https://web.archive.org/web/20171019074321/http://faseb.org/Science-Policy--Advocacy-and-Communications/Federal-Funding-Data/NIH-Research-Funding-Trends.aspx

## REFERENCES

1. Zhulin I.B. (2015) Databases for Microbiologists. J Bacteriol., 197, 2458–67. doi: 10.1128/JB.00330-15

2. Wren J.D. (2016) Bioinformatics programs are 31-fold over-represented among the highest impact scientific papers of the past two decades. Bioinformatics, 32, 2686–91. doi: 10.1093/bioinformatics/btw284

3. Fernández-Suárez X.M., Galperin M.Y. (2013) The 2013 Nucleic Acids Research Database Issue and the online Molecular Biology Database Collection. Nucleic Acids Res., 41, D1–7. doi: 10.1093/nar/gks1297

4. Imker H.J. (2018) 25 Years of Molecular Biology Databases: A Study of Proliferation, Impact, and Maintenance. Front. Res. Metr. Anal., 3, 18. doi: 10.3389/frma.2018.00018

5. Kalumbi D., Ellis L.B.M. (1998) The demise of public data on the web? Nature Biotechnology, 16,1323–1324.

6. Kaiser J. (2016) Funding for key data resources in jeopardy. Science, 351, 14. doi: 10.1126/science.351.6268.14

7. Anderson W.P. (2017) Data management: A global coalition to sustain core data. Nature, 543, 179–179. doi: 10.1038/543179a

8. Anderson W., Apweiler R., Bateman A., Bauer G.A., Berman H., Blake J.A., et al. (2017) Towards Coordinated International Support of Core Data Resources for the Life Sciences. bioRxiv, 110825. doi: 10.1101/110825

9. Galperin M.Y., Fernández-Suárez X.M., Rigden D.J. (2017) The 24th annual Nucleic Acids Research database issue: a look back and upcoming changes. Nucleic Acids Res., 45, 5627–5627. doi: 10.1093/nar/gkx021

10. Imker H. (2018) Molecular Biology Databases Published in Nucleic Acids Research between 1991-2016. University of Illinois at Urbana-Champaign, doi: 10.13012/B2IDB-4311325_V1

11. Grassano N., Rotolo D., Hutton J., Lang F., Hopkins M.M. (2016) Funding Data from Publication Acknowledgments: Coverage, Uses, and Limitations. J Assoc Inf Sci Technol., 68, 999–1017. doi: 10.1002/asi.23737

12. Viergever R.F., Hendriks T.C.C. The 10 largest public and philanthropic funders of health research in the world: what they fund and how they distribute their funds. Health Res Policy Syst., 2016, 14, 2. doi: 10.1186/s12961-015-0074-z

13. Maron N.L. (2014) A Guide to the Best Revenue Models and Funding Sources for your Digital Resources. Ithaka S+C and JISC, Available from: http://sr.ithaka.org/?p=22805

14. Kitchin R., Collins S., Frost D. (2015) Funding Models for Open Access Repositories Maynooth: Maynooth University and Dublin: the Royal Irish Academy and Trinity College Dublin. doi: 10.3318/DRI.2015.4

15. Gabella C., Durinx C., Appel R. (2018) Funding knowledgebases: Towards a sustainable funding model for the UniProt use case. F1000Research, 6, 2051. doi: 10.12688/f1000research.12989.2

16. Wickham H. (2017) tidyverse: Easily Install and Load “Tidyverse” Packages. Available from: https://CRAN.R-pro-ject.org/package=tidyverse

17. Komsta L., Novomestky F. (2015) moments: Moments, cumulants, skewness, kurtosis and related tests. Available from: https://CRAN.R-project.org/package=moments

18. Wickham H. (2009) ggplot2: Elegant Graphics for Data Analysis. Springer-Verlag New York, Available from: http://ggplot2.org

19. OECD. (2017) Business models for sustainable research data repositories. Available from: http://www.oecd-ili-brary.org/science-and-technology/business-models-for-sustainable-research-data-repositories_302b12bb-en

20. Ember C., Hanisch R. (2013) Sustaining Domain Repositories for Digital Data: A White Paper. Available from: http://datacommunity.icpsr.umich.edu/sites/default/files/WhitePaper_ICPSR_SDRDD_121113.pdf

21. Bourne P.E., Lorsch J.R., Green E.D. (2015) Perspective: Sustaining the big-data ecosystem. Nature, 527, S16–7. doi: 10.1038/527S16a

22. Merali Z., Giles J. (2005) Databases in peril. Nature, 435, 1010–1. doi: 10.1038/4351010a

23. Baker M. (2012) Databases fight funding cuts. Nature, 489, 19. doi:10.1038/489019a

24. Attwood T.K., Agit B., Ellis L.B.M. (2015) Longevity of Biological Databases. EMBnet.journal., 21, e803. doi: 10.14806/ej.21.0.803

25. Berman F., Cerf V. (2013) Who Will Pay for Public Access to Research Data? Science, 341, 616–7. doi: 10.1126/science.1241625

26. Oliver S.G., Lock A., Harris M.A., Nurse P., Wood V. (2016) Model organism databases: essential resources that need the support of both funders and users. BMC Biol., 22, 49. doi: 10.1186/s12915-016-0276-z

27. Kirlew P.W. (2011) Life Science Data Repositories in the Publications of Scientists and Librarians. Issues Sci Technol Librarianship, Spring, 65. doi: 10.5062/F4X63JT2

