## Supplemental Tables for "Who Bears the Burden of Long-Lived Molecular Biology Databases?"

### Supplemental Materials

S1. Individual funding organizations

S2. Individual databases

S3. Individual operating organizations

**Table S1.** Individual funding organizations identified as funding databases in *NAR* Database Issue articles. Organizational codes are A (academic), C (consortium/collective), G (government), I (industry), P (philanthropic), and S (society/association.

| | FUNDER | ORG CODE | DB COUNT |
|---|---|---|---|
| 1 | US National Institutes of Health | G | 29 |
| 2 | European Commission | G | 13 |
| 3 | UK Biotechnology and Biological Sciences Research Council | G | 7 |
| 4 | US Department of Energy | G | 7 |
| 5 | Ministry of Education Culture Sports Science and Technology of Japan | G | 6 |
| 6 | US National Science Foundation | G | 6 |
| 7 | Japan Science and Technology Agency | G | 5 |
| 8 | European Molecular Biology Laboratory | G | 5 |
| 9 | Danish Ministry of Higher Education and Science | G | 5 |
| 10 | UK Medical Research Council | G | 4 |
| 11 | Wellcome Trust | P | 3 |
| 12 | Kazusa DNA Research Institute Foundation | P | 3 |
| 13 | University of Tokyo | A | 2 |
| 14 | Aarhus University | A | 2 |
| 15 | SWEGENE Consortium | C | 2 |
| 16 | Swiss National Science Foundation | G | 2 |
| 17 | Agence Nationale de la Recherche | G | 2 |
| 18 | Howard Hughes Medical Institute | P | 2 |
| 19 | Adam Mickiewicz University | A | 1 |
| 20 | Kyoto University | A | 1 |
| 21 | Poitiers University Foundation | A | 1 |
| 22 | University of Leicester | A | 1 |
| 23 | Universite de Montpellier | A | 1 |
| 24 | Oxford University Press | A | 1 |
| 25 | University of California Irving | A | 1 |
| 26 | Johns Hopkins University | A | 1 |
| 27 | University of Manchester | A | 1 |
| 28 | Universidad Nacional Autonoma de Mexico | A | 1 |
| 29 | Cornell University | A | 1 |
| 30 | Dutch Working Group of Tumor Cytogenetics | C | 1 |
| 31 | Cancer Cytogenomics Microarray Consortium US | C | 1 |
| 32 | National Centre for Research on Growth and Development | C | 1 |
| 33 | BioCampus Montpellier | C | 1 |
| 34 | Grand Plateau Technique pour la Recherche | C | 1 |
| 35 | Ciber Enfermedades Raras | C | 1 |
| 36 | King Abdelaziz City for Science and Technology | G | 1 |
| 37 | French Ministry of Higher Education and Scientific Research | G | 1 |
| 38 | Czech Science Foundation | G | 1 |
| 39 | Polish National Science Centre | G | 1 |
| 40 | Health Research Council of New Zealand | G | 1 |
| 41 | Centre National de la Recherche Scientifique | G | 1 |
| 42 | Ministere de l Enseignement Superieur et de la Recherche | G | 1 |
| 43 | Spanish Ministry of Economy Industry and Competitiveness | G | 1 |
| 44 | Diputacion General de Aragon | G | 1 |
| 45 | Polish Ministry of Science and Higher Education | G | 1 |
| 46 | Engineering and Physical Sciences Research Council | G | 1 |
| 47 | Swiss Federal Office of Education and Science | G | 1 |
| 48 | US Department of Agriculture | G | 1 |
| 49 | Science Foundation Ireland | G | 1 |
| 50 | German Federal Ministry of Education and Research | G | 1 |
| 51 | Italian Ministry of Education Universities and Research | G | 1 |
| 52 | Italian National Research Council | G | 1 |
| 53 | Histogenetics | I | 1 |
| 54 | One Lambda Inc | I | 1 |
| 55 | Conexio | I | 1 |
| 56 | Abbott Molecular Laboratories Inc | I | 1 |
| 57 | GenProbe | I | 1 |
| 58 | LabCorp | I | 1 |
| 59 | Life Technologies | I | 1 |
| 60 | Olersup SSP | I | 1 |
| 61 | 454 Sequencing | I | 1 |
| 62 | BAG Healthcare | I | 1 |
| 63 | Innotrain Diagnostik GMBH | I | 1 |
| 64 | Lundbeck Foundation | I | 1 |
| 65 | GlaxoSmithKline | I | 1 |
| 66 | BioFocus | I | 1 |
| 67 | New England Biolabs | I | 1 |
| 68 | Isis Pharmaceuticals | I | 1 |
| 69 | Microsoft | I | 1 |
| 70 | BioRegioN GmbH | I | 1 |
| 71 | BIOBASE GmbH | I | 1 |
| 72 | Anthony Nolan | P | 1 |
| 73 | Be the Match Foundation | P | 1 |
| 74 | Deutsche Knochenmarkspenderdatei | P | 1 |
| 75 | Rose and Zentrum Knochenmarkspender Register Deutschland | P | 1 |
| 76 | Imperial Cancer Research Fund | P | 1 |
| 77 | Child Cancer Foundation | P | 1 |
| 78 | Ellison Foundation | P | 1 |
| 79 | Sandler Family Supporting Foundation | P | 1 |
| 80 | National Foundation for Cancer Research | P | 1 |
| 81 | Association des Cytogeneticiens de Langue Francaise | S | 1 |
| 82 | Belgian Society of Human Genetics | S | 1 |
| 83 | French National League against Cancer | S | 1 |
| 84 | Berufsverband Deutscher Humangenetiker | S | 1 |
| 85 | European Federation for Immunogenetics | S | 1 |
| 86 | American Society for Histocompatibility and Immunogenetics | S | 1 |
| 87 | AsiaPacific Histocompatibility and Immunogetics Association | S | 1 |
| 88 | Fonds der Chemischen Industrie eV | S | 1 |
| 89 | Royal Society | S | 1 |

**Table S2.** Funding organizations identified for individual databases. Organizational codes are A (academic), C (consortium/collective), G (government), I (industry), P (philanthropic), and S (society/association). Database identifiers (DB ID) are as reported in (Imker, 2018).

| | DB ID | NAME | NAR DEBUT YEAR | ALL FUNDER ORG CODES | CONDENSED ORG CODES |
|---|---|---|---|---|---|
| 1 | MBDC0001 | DDBJ: DNA Data Bank of Japan | 1997 | GG | G |
| 2 | MBDC0002 | EMBL-EBI ENA: European Nucleotide Archive | 1991 | GGG | G |
| 3 | MBDC0003 | NCBI Genbank | 1991 | G | G |
| 4 | MBDC0007 | NCBI COG: Clusters of Orthologous Groups | 2000 | G | G |
| 5 | MBDC0015 | GXD: mouse Gene Expression Database | 1999 | GG | G |
| 6 | MBDC0021 | RECODE | 2001 | GG | G |
| 7 | MBDC0030 | CUTG: Codon Usage Tabulated from GenBank | 1991 | GP | GP |
| 8 | MBDC0031 | DBTBS: Database of Transcription in B. subtilis | 2001 | AGG | AG |
| 9 | MBDC0033 | EPD: Eukaryotic Promoter Database | 1998 | G | G |
| 10 | MBDC0043 | rrnDB | 2001 | GGG | G |
| 11 | MBDC0051 | WormBase | 2001 | GGGP | GP |
| 12 | MBDC0068 | CyanoBase | 1998 | GGP | GP |
| 13 | MBDC0069 | EcoGene | 2000 | G | G |
| 14 | MBDC0071 | FlyBase | 1994 | GG | G |
| 15 | MBDC0072 | DB-AT: Database of Apicomplexan Transcriptomes | 2001 | GG | G |
| 16 | MBDC0075 | JGI GOLD: Genomes OnLine Database | 2001 | G | G |
| 17 | MBDC0084 | MITOMAP | 1996 | ACGGGGP | ACGP |
| 18 | MBDC0087 | MGD: Mouse Genome Database | 1997 | G | G |
| 19 | MBDC0091 | PlasmoDB | 2001 | G | G |
| 20 | MBDC0097 | TAIR: The Arabidopsis Information Resource | 2001 | AGGI | AGI |
| 21 | MBDC0101 | ZFIN: Zebrafish Information Network | 2001 | G | G |
| 22 | MBDC0108 | EcoCyc | 1996 | G | G |
| 23 | MBDC0112 | KEGG: Kyoto Encyclopedia of Genes and … | 1999 | G | G |
| 24 | MBDC0116 | RegulonDB | 1998 | AG | AG |
| 25 | MBDC0119 | ALFRED: Allele Frequency Database | 2000 | G | G |
| 26 | MBDC0122 | Atlas of Genetics and Cytogenetics in Oncology … | 1999 | ACCGGSSSS | ACGS |
| 27 | MBDC0131 | GWAS Centeral | 2000 | AGI | AGI |
| 28 | MBDC0143 | OMIM: Online Mendelian Inheritance in Man | 1994 | AG | AG |
| 29 | MBDC0152 | MTB: Mouse Tumor Biology Database | 1999 | G | G |
| 30 | MBDC0162 | ESTHER: ESTerases, alpha-beta Hydrolase … | 1996 | G | G |
| 31 | MBDC0166 | GPCRDB: G Protein-Coupled Receptor Database | 1998 | GGGI | GI |
| 32 | MBDC0171 | HUGE | 1999 | P | P |
| 33 | MBDC0172 | IMGT: ImMunoGeneTics database | 1997 | ACCGGGG | ACG |
| 34 | MBDC0173 | EMBL-EBI IPD IMGT/HLA | 2001 | GIIIIIIIIIIPPPPPSSS | GIPS |
| 35 | MBDC0177 | MEROPS | 1999 | P | P |
| 36 | MBDC0194 | PIR: Protein Identification Resouce | 1991 | G | G |
| 37 | MBDC0195 | Ribonuclease P Database | 1994 | GI | GI |
| 38 | MBDC0199 | TIGRFAMs | 2001 | G | G |
| 39 | MBDC0207 | EMBL-EBI InterPro | 2001 | GG | G |
| 40 | MBDC0210 | Pfam | 1998 | GGPP | GP |
| 41 | MBDC0212 | PRINTS | 1994 | AGGGGIS | AGIS |
| 42 | MBDC0215 | PROSITE | 1991 | GG | G |
| 43 | MBDC0218 | SMART: Simple Modular Architecture Resea… | 1999 | G | G |
| 44 | MBDC0221 | AAIndex: Amino Acid Index Database | 1999 | AAGG | AG |
| 45 | MBDC0224 | REBASE: Restriction Enzymes and methylases … | 1991 | I | I |
| 46 | MBDC0230 | 5S rRNA Data Bank (Erdmann) | 1991 | A | A |
| 47 | MBDC0232 | ARED: Adenylate uridylate-Rich Elements … | 2001 | G | G |
| 48 | MBDC0243 | RDP: Ribosomal Database Project | 1991 | GGGG | G |
| 49 | MBDC0248 | SRPDB: Signal Recognition Particle Database | 1992 | ACGGG | ACG |
| 50 | MBDC0250 | tmRDB: tmRNA database | 1998 | ACGGG | ACG |
| 51 | MBDC0252 | UTRdb: UnTranslated Regions of eukaryotic mR … | 1998 | GG | G |
| 52 | MBDC0258 | CATH: Class, Architecture, Topology, Homology | 1999 | GGP | GP |
| 53 | MBDC0270 | NCBI MMDB: Molecular Modeling DataBase | 1999 | G | G |
| 54 | MBDC0276 | RCSB PDB: Protein Data Bank | 2000 | GGG | G |
| 55 | MBDC0278 | EMBL-EBI PDBsum | 2001 | GGGGG | G |
| 56 | MBDC0282 | SCOP: Structural Classification of Proteins | 1997 | GG | G |
| 57 | MBDC0294 | IGC: Imprinted Gene Catalogue | 2001 | CGP | CGP |
| 58 | MBDC0331 | MODBASE | 2000 | GGGP | GP |
| 59 | MBDC0340 | TRANSFAC | 1996 | GGII | GI |
| 60 | MBDC0357 | SGD: Saccharomyces Genome Database | 1998 | G | G |
| 61 | MBDC0381 | ncRNAdb: Noncoding RNAs database | 1999 | GPS | GPS |
| 62 | NAR9161 | MBDC: Molecular Biology Database Collection | 1999 | AG | AG |
| 63 | NAR9294 | tmRNA Website | 1998 | G | G |

**Table S3.** Individual operating organizations identified. Organizational codes are A (academic), C (consortium/collective), G (government), I (industry), P (philanthropic), and S (society/association).

| | HOST | ORG CODE | DB COUNT | TOTAL DB COUNT / OPER ORG CODE |
|---|---|---|---|---|
| **1** | European Molecular Biology Laboratory | G | 8 | 10 |
| **2** | Swiss Institute of Bioinformatics | G | 4 | 10 |
| **3** | US National Institutes of Health | G | 4 | 10 |
| **4** | Kazusa DNA Research Institute | P | 3 | 5 |
| **5** | Jackson Laboratory | P | 3 | 5 |
| **6** | Kyoto University | A | 2 | 28 |
| **7** | University of Tokyo | A | 2 | 28 |
| **8** | Center for NonCoding RNA in Technology and Health | C | 2 | 4 |
| **9** | Adam Mickiewicz University | A | 1 | 28 |
| **10** | Yale University | A | 1 | 28 |
| **11** | University College London | A | 1 | 28 |
| **12** | University of Miami | A | 1 | 28 |
| **13** | University of Copenhagen | A | 1 | 28 |
| **14** | University of Leicester | A | 1 | 28 |
| **15** | University of Otago | A | 1 | 28 |
| **16** | Universite de Montpellier | A | 1 | 28 |
| **17** | Oxford University Press | A | 1 | 28 |
| **18** | University of California Irving | A | 1 | 28 |
| **19** | University of California San Francisco | A | 1 | 28 |
| **20** | Johns Hopkins University | A | 1 | 28 |
| **21** | University of Delaware | A | 1 | 28 |
| **22** | Georgetown University Medical Center | A | 1 | 28 |
| **23** | University of Manchester | A | 1 | 28 |
| **24** | Rutgers University | A | 1 | 28 |
| **25** | The State University of New Jersey | A | 1 | 28 |
| **26** | University of California San Diego | A | 1 | 28 |
| **27** | Michigan State University | A | 1 | 28 |
| **28** | University College Cork | A | 1 | 28 |
| **29** | National Autonomous University of Mexico | A | 1 | 28 |
| **30** | North Carolina State University | A | 1 | 28 |
| **31** | University of Michigan | A | 1 | 28 |
| **32** | Stanford University | A | 1 | 28 |
| **33** | University of Bari | A | 1 | 28 |
| **34** | University of Oregon | A | 1 | 28 |
| **35** | King Faisal Specialist Hospital and Research Center | G | 1 | 10 |
| **36** | Institute of Scientific and Technical Information | G | 1 | 10 |
| **37** | DDJB Center | G | 1 | 10 |
| **38** | French National Institute for Agricultural Research | G | 1 | 10 |
| **39** | DOE Joint Genome Institute | G | 1 | 10 |
| **40** | MRC Laboratory of Molecular Biology | G | 1 | 10 |
| **41** | Sandia National Laboratories | G | 1 | 10 |
| **42** | SRI International | P | 1 | 5 |
| **43** | Phoenix Bioinformatics Corporation | P | 1 | 5 |
| **44** | J Craig Venter Institute | P | 1 | 5 |
| **45** | FLyBase Consortium | C | 1 | 4 |
| **46** | EuPathDB Consortium | C | 1 | 4 |
| **47** | Wormbase Consortium | C | 1 | 4 |
| **48** | New England Biolabs | I | 1 | 2 |
| **49** | GeneXplain | I | 1 | 2 |
| **50** | Institute of Bioorganic Chemistry Polish Academy of Sciences | S | 1 | 1 |
